# GeneVector: Identification of transcriptional programs using dense vector representations defined by mutual information

**DOI:** 10.1101/2022.04.22.487554

**Authors:** Nicholas Ceglia, Zachary Sethna, Samuel S. Freeman, Florian Uhlitz, Viktoria Bojilova, Nicole Rusk, Bharat Burman, Andrew Chow, Sohrab Salehi, Farhia Kabeer, Samuel Aparicio, Benjamin Greenbaum, Sohrab P. Shah, Andrew McPherson

**Author notes:** Correspondence, 1275 York Ave, New York, NY 10065. Authors contributed equally (Zachary Sethna, Samuel S. Freeman).

## Abstract

Deciphering individual cell phenotypes from cell-specific transcriptional processes requires high dimensional single cell RNA sequencing. However, current dimensionality reduction methods aggregate sparse gene information across cells, without directly measuring the relationships that exist between genes. By performing dimensionality reduction with respect to gene co-expression, low-dimensional features can model these gene-specific relationships and leverage shared signal to overcome sparsity. We describe GeneVector, a scalable framework for dimensionality reduction implemented as a vector space model using mutual information between gene expression. Unlike other methods, including principal component analysis and variational autoencoders, GeneVector uses latent space arithmetic in a lower dimensional gene embedding to identify transcriptional programs and classify cell types. In this work, we show in four single cell RNA-seq datasets that GeneVector was able to capture phenotype-specific pathways, perform batch effect correction, interactively annotate cell types, and identify pathway variation with treatment over time.

## Introduction

Maintenance of cell state and execution of cellular function are based on coordinated activity within networks of related genes. To approximate these connections, transcriptomic studies have conceptually organized the transcriptome into sets of co-regulated genes, termed *gene programs* or *metagenes* (J. M. Stuart 2003; Svensson et al. 2020). The first intuitive step to identify such co-regulated genes is the reduction of dimensionality for sparse expression measurements: high dimensional gene expression data is compressed into a minimal set of explanatory features that highlight similarities in cellular function. However, to map existing biological knowledge to each cell, the derived features must be interpretable at the gene level.

To find similarities in lower dimensions, biology can borrow from the field of natural language processing (NLP). NLP commonly uses dimensionality reduction to identify word associations within a body of text (Mikolov et al. 2013, Pennington, Socher, and Manning 2014). To find contextually similar words, NLP methods make use of vector space models to represent similarities in a lower dimensional space. Similar methodology has been applied to bulk RNA-seq expression for finding co-expression patterns (Du et al. 2019). Inspired by such work, we developed a tool that generates gene vectors based on single cell RNA (scRNA)-seq expression data. While current methods reduce dimensionality with respect to sparse expression across each cell, our tool produces a lower dimensional embedding with respect to each gene. The vectors derived from GeneVector provide a framework for identifying metagenes within a gene co-expression graph and relating these metagenes back to each cell using latent space arithmetic.

The most pervasive method for identifying the sources of variation in scRNA-seq studies is principal component analysis (PCA) (Hao et al. 2021; Wolf, Angerer, and Theis 2018; Zheng et al. 2017). The relationship of principal components to gene expression is linear, allowing lower dimensional structure to be directly related to variation in expression. A PCA embedding is an ideal input for building a nearest neighbor graph for unsupervised clustering algorithms (Traag, Waltman, and van Eck 2019) and visualization methods including t-SNE (Pezzotti et al. 2017) and UMAP (McInnes et al. 2018). However, the assumption of a continuous multivariate gaussian distribution creates distortion in modeling read counts generated by a true distribution that is over-dispersed, possibly zero-inflated (Jiang et al. 2022), with positive support and mean close to 0 (Svensson et al. 2020). Despite such issues, gene programs generated from PCA loadings have been used to generate metagenes that explain each principal component (Tsuyuzaki et al. 2020). While these loadings highlight sets of genes that explain each orthogonal axis of variation, pathways and cell type signatures can be conflated within a single axis of variation.

In addition to PCA, more sophisticated methods have been developed to better handle the specific challenges of scRNA data. The single cell variational inference (scVI) framework generates an embedding using non-linear autoencoders that can be used in a range of analyses including normalization, batch correction, gene-dropout correction, and visualization (Lopez et al. 2018). While scVI embeddings show improved performance over traditional PCA-based analysis in these tasks, they have a non-linear relationship to the original count matrix that may distort the link between structure in the generated embedding and potentially identifiable gene programs (Svensson et al. 2020). A subsequent method uses a linearly decoded variational autoencoder (LDVAE), which combines a Variational Autoencoder with a factor model of negative binomial distributed read counts to learn an interpretable linear imbedding of cell expression profiles (Svensson et al. 2020). However, the relationship between gene expression and cell representation is still tied to correlated variation across cells, which may confound co-varying pathway and phenotypic signatures.

Recognizing the importance of modeling the non-linearity of gene expression and the complexity of statistical dependencies between genes, several methods have adopted information theoretical approaches. Many of these methods use mutual information (MI), an information theoretic measure of the statistical dependence between two variables. ARACNE (Margolin et al. 2006) uses MI to prune independent and indirectly interacting genes during construction of a gene regulatory network from microarray expression data. PIDC (Chan, Stumpf, and Babtie 2017) identifies regulatory relationships using partial information decomposition (PID), a measure of the dependence between triples of variables. The authors apply PIDC to relatively high-depth single-cell qPCR datasets and restrict their analysis to on the order of hundreds of genes. More recently, IQCELL (Heydari et al. 2022) uses MAGIC (van Dijk et al. 2018) to impute missing gene expression, builds a GRN from pairwise MI between genes, and applies a series of filters to produce a GRN composed of only functional relationships. The authors use IQCELL to identify known causal gene interactions in scRNA-seq data from mouse T-cell and red blood cell development experiments. Because of the success of these methods, we hypothesized that MI could be combined with vector space models to produce a meaningful low dimensional representation of genes from scRNA data.

In this work, we present GeneVector (**Figure 1**) as a framework for generating low dimensional embeddings constructed from the mutual information between genes. GeneVector summarizes coexpression of genes as mutual information between the probability distribution of read counts across cells. We showcase GeneVector on 4 scRNA datasets produced from a diverse set of experiments: peripheral blood mononuclear cells (PBMCs) subjected to interferon beta stimulation (Kang et al. 2018), the Tumor Immune Cell Atlas (TICA) (Nieto et al. 2021), treatment naive multi-site samples from High Grade Serous Ovarian Cancers (HGSOC) (Vázquez-García et al. 2022) and a time series of cisplatin treatment in patient-derived xenografts (PDX) of triple negative breast cancer (TNBC) (Salehi et al. 2021). We first confirm GeneVector’s ability to identify putatively co-regulated gene pairs from sparse single cell expression measurements using the TICA and PBMC datasets. Next, we show that GeneVector can identify metagenes corresponding to cell-specific transcriptional processes in PBMCs, including interferon activated gene expression. We further show that latent space arithmetic can be used to accurately label cell types in the TICA dataset and validate our cell type predictions against published annotations. We use vector space arithmetic to directly map metagenes to site specific changes in primary and metastatic sites in the HGSOC dataset, capturing changes in MHC class I expression and epithelial-mesenchymal transition (EMT). Finally, we show GeneVector can identify cisplatin treatment dependent transcriptional programs related to TGF-beta in TNBC PDXs.

**Figure 1:**
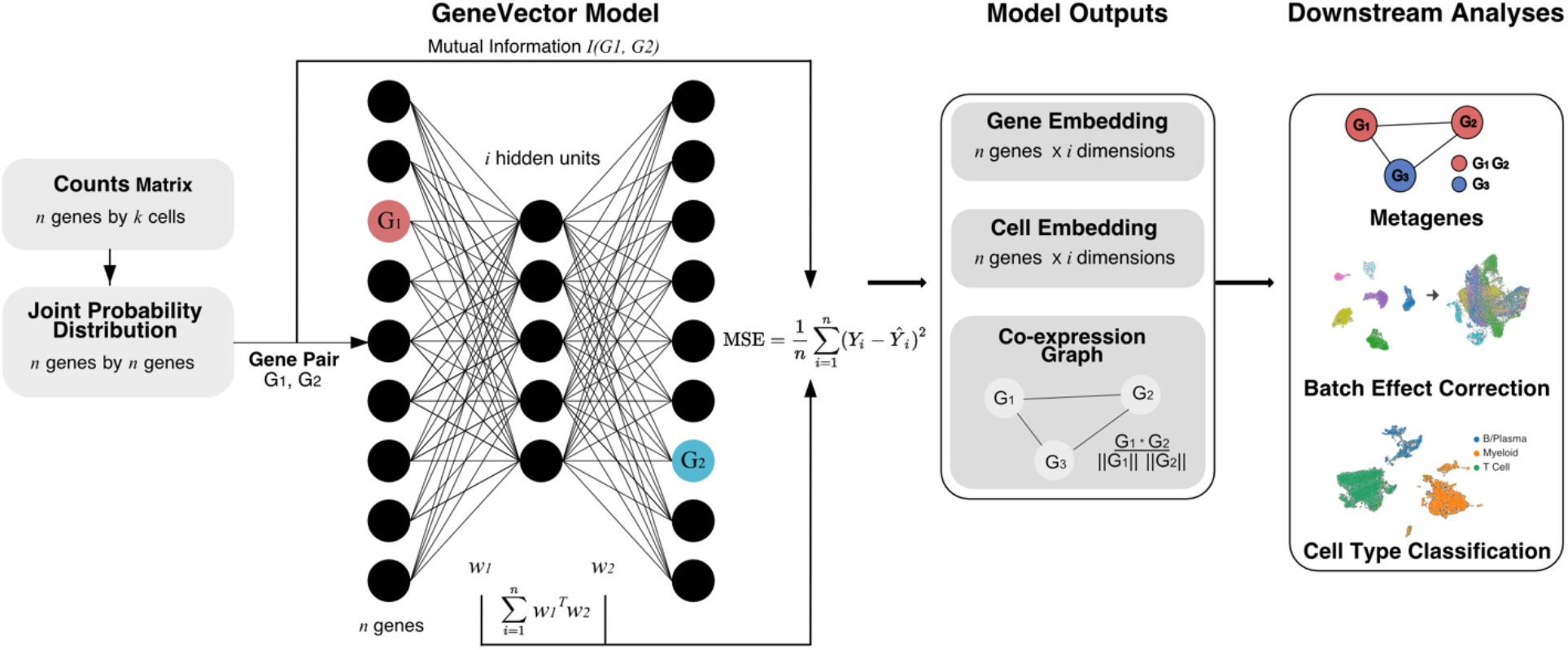
GeneVector Framework. Overview of GeneVector framework starting from single cell read counts. Mutual information is computed on the joint probability distribution of read counts for each gene pair. Each pair is used to train a single layer neural network where the MSE loss is evaluated from the model output (w_1_^T^w_2_) with the mutual information between genes. From the resulting weight matrix, a gene embedding, cell embedding, and co-expression similarity graph are constructed. Using vector space arithmetic, downstream analyses include identification of cell-specific metagenes, batch effect correction, and cell type classification.

## Results

### Defining the GeneVector Framework

We trained a single layer neural network over all gene pairs to generate low dimensional gene embeddings and identify metagenes from a co-expression similarity graph. The input weights (*w_1_*) and output weights (*w_2_*) are updated with adaptive gradient descent (ADADELTA) (Zeiler 2012). Gene coexpression relationships are defined using mutual information (**Methods:** Mutual Information) computed from a joint probability distribution of expression counts. Training loss is evaluated as the mean squared error of mutual information with the model output, defined as *w_1_^T^w_2_.* The final latent space is a matrix defined as a series of vectors for each gene.

Gene vectors produced by the framework are useful for several fundamental gene expression analyses. Gene vectors weighted by expression in each cell are combined to generate the cell embedding analysis of cell populations and their relationships to experimental covariates. The cell embedding can be batch corrected by using vector arithmetic to identify vectors that represent batch effects and then shift cells in the opposite direction (**Methods:** Batch correction). A co-expression graph is constructed in which each node is defined as a gene and each edge is weighted by cosine similarity. After generating the coexpression graph, we use Leiden clustering (Traag, Waltman, and van Eck 2019) to identify metagene clusters. Further downstream analysis of the cell embedding includes phenotype assignment based on sets of marker genes and computation of the distances between cells and metagenes to highlight changes related to experimental covariates (**Figure 1**).

To perform cell type assignment, a set of known marker genes is used to generate a representative vector for each cell type, where each gene vector is weighted by the normalized gene expression. The cosine similarity of each possible phenotype is computed between the cell vector and the marker gene vector. SoftMax is applied to cosine distances to obtain a pseudo-probability over each phenotype (**Methods**: Cell type assignment). Discrete labels can be assigned to cells by selecting the phenotype corresponding to the maximum pseudo-probability. More generally, gene vectors can be composed together to describe novel gene expression features. Cell or gene vectors can then be compared against these feature vectors to evaluate the relevance of that feature to a given cell or gene (**Methods**: Generation of Predictive Genes).

### Robust Inference of Gene Co-Regulation with GeneVector

Our model relies on the advantages of mutual information to define relationships between genes, as opposed to other distance metrics. To evaluate how MI contributes to the observed performance of the model, we first validated that the vectors inferred by GeneVector capture semantic qualities of genes including pathway memberships and regulatory relationships. Specifically, we assessed the extent to which pairs of genes within the same pathway, or expressed within the same cell types, produced similar vector representations in the PBMC dataset. As a ground truth we computed, for each gene pair, the number of combined pathways from Reactome and MSigDB cell type signatures (C8) (Gillespie et al. 2022, Liberzon et al. 2011; Subramanian et al. 2005) for which both genes were members. In addition to training GeneVector with an MI target, we trained on Pearson correlation coefficient to evaluate the relative benefit of MI on the accuracy of the model output. For both the correlation and MI based models, cosine similarities between gene vectors showed a stronger relationship with the number of shared pathways and cell type signatures than randomly shuffled gene pairs (**Figure 2A,C**). Comparing the MI model and the correlation model directly, the MI model produced a much stronger relationship than the correlation based model (r^2^ = 0.233 vs. r^2^ = 0.093, **Figure 2B,D**). To provide a pathway specific example, we found that the most similar genes by cosine similarity to IFIT1 (a known interferon stimulated gene) using the correlation objective were less coherent in terms of ISG pathway membership signal (6 of 16 genes were in the Interferon Signaling Reactome pathway R-HSA-913531) (**Figure 2E**) than with mutual information (12 of 16 genes) (**Figure 2F**). The greater proportion of interferon stimulated pathway genes with high similarity to IFIT1 using a mutual information objective function (**Figure 2F vs. 2E**) is consistent with the improved correlation between pathway co-membership and cosine similarity over the Pearson correlation coefficient objective (**Figure 2D vs. 2B**).

**Figure 2:**
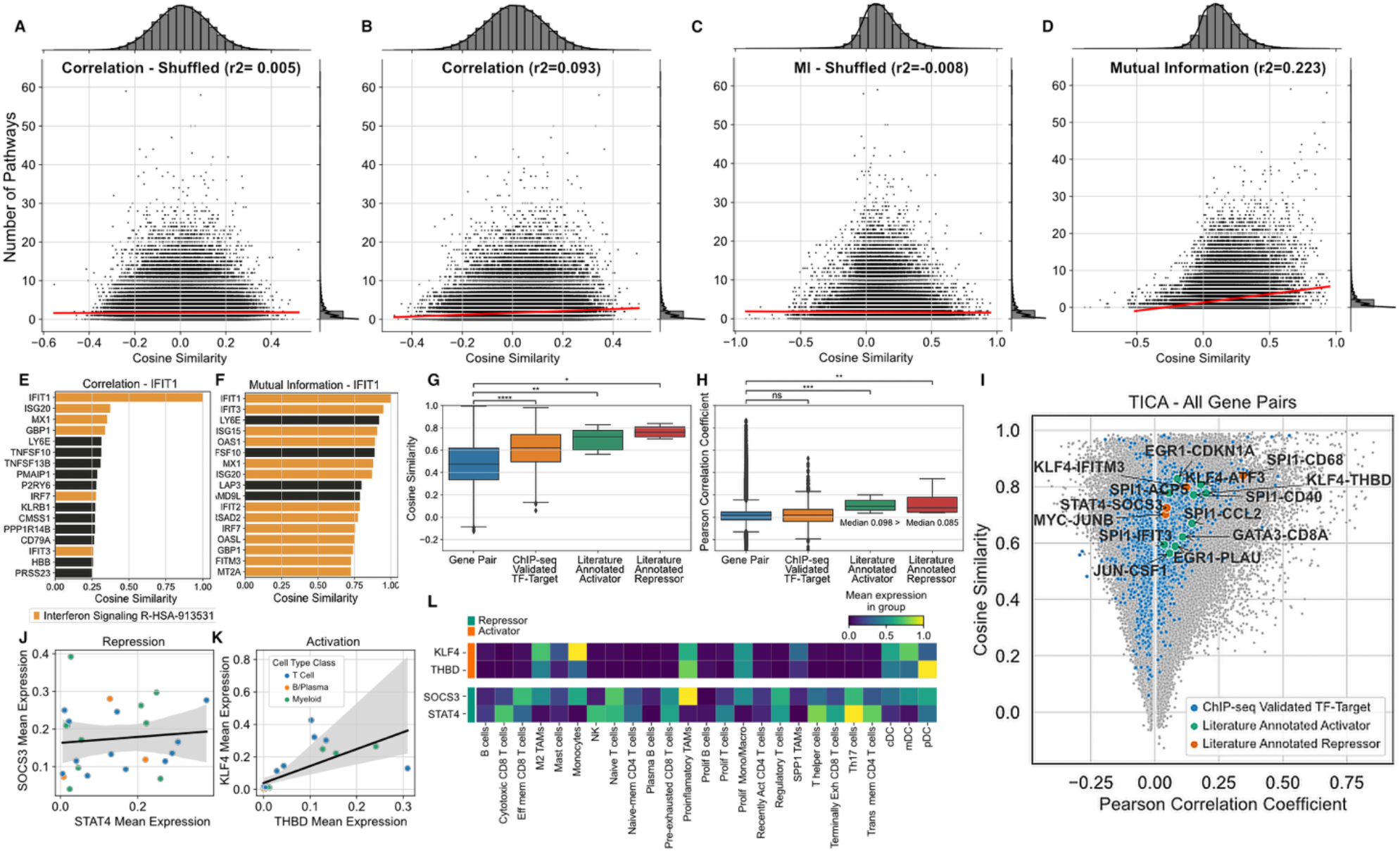
Comparison of Results using Mutual Information. A-D) Pathway co-membership vs. cosine similarity between gene vectors for all gene pairs in PBMCs. Each point represents one gene pair, and plots show the number of pathways (combined Reactome and MSigDB cell type signatures [C8]) that contain both genes (y-axis) and the cosine distance between the two genes (x-axis). The results show both correlation (A,B) and MI (C,D) based GeneVector. In addition to a standard set of results (B, D), a baseline relationship between pathway co-membership and cosine similarity is established by performing an identical analysis over randomly shuffled gene (A, C). E) Top 16 most similar genes by cosine similarity to IFIT1 using correlation coefficient. Genes in the interferon signaling pathway are colored orange. F) Top 16 most similar genes to IFIT1 after training GeneVector using mutual information shows a higher number of interferon signaling pathway genes. G) Cosine similarity for un-annotated gene pairs, ChIP-Seq annotated TF-targets pairs, and literature annotated activator or repressor TF-target pairs. H) Pearson correlation coefficient for un-annotated gene pairs, ChIP-Seq annotated TF-targets pairs, and literature annotated activator or repressor TF-target pairs. I) Cosine similarity versus correlation coefficient for gene pairs in the TICA dataset with TF-target gene pairs highlighted (blue) and colored by activator/repressor status (green/orange respectively). J-K) Linear regression of mean expression per cell type for repressor TF-target pair SOCS3-STAT4 and activator TF-target pair KLF-THBD, respectively. L) Mean expression for SOCS3-STAT4 and KLF4-THBD across annotated cell types. (***p < 0.001, **p < 0.01, *p < 0.05, ns=not significant)

Next, we sought to understand whether GeneVector was able to capture relationships between genes with known interactions such as transcription factors (TF) and their targets. GeneVector cosine similarities of TF-target pairs annotated based on ChIP-Seq data (Q. Zhang et al. 2020) were significantly increased relative to un-annotated gene pairs in the TICA dataset (**Figure 2G**). We also considered literature annotated activator-target and repressor-target TF-target gene pairs (Han et al. 2018). As expected, activator-target pairs showed increased cosine similarity above unannotated pairs. Importantly, repressor-target pairs showed an equally strong increase in cosine similarity above unannotated pairs highlighting GeneVector’s ability to identify a diversity of statistical dependencies between co-regulated genes. By comparison, Pearson correlation coefficients of ChIP-seq annotated TF-target pairs were not significantly different from unannotated gene pairs (**Figure 2H**). Both activator-target and repressor-target pairs were significantly different from unannotated pairs, though correlation was on-average positive for both repressor-target and activator-target pairs (**Figure 2H**). In fact, while many repressor-target pairs had high cosine similarity indicative of a meaningful regulatory relationship, their correlation coefficients were always positive (**Figure 2I**). For example, SOCS3-STAT4 exhibited the lowest correlation of annotated repressor-target pairs (r^2^=0.017) and aggregating expression across cell types showed an absence of any relationship between these genes (**Figure 2J**). In contrast, analysis of activator-target pair KLF4-THBD revealed a positive correlation driven by co-expression in myeloid cells and T cells (**Figure 2K**). This relationship is further evidenced when looking at the expression in more detailed cell type annotations (**Figure 2L**). Identification of mutually exclusive expression is a theoretical benefit of correlation-based similarity measures, however, the sparsity of scRNA likely results in positive or low correlation even for known examples of mutual exclusivity. In summary, the vector space produced by GeneVector successfully recovers the latent similarities between functionally related genes, including negative regulators and their targets, overcoming the sparsity of scRNA data that confounds simpler approaches.

### Fast and Accurate Cell Type Classification using GeneVector

Comparative analysis of gene expression programs across large cohorts of patients can potentially identify transcriptional patterns in common cell types shared between many cancers. However, classification of cell types using methods such as CellAssign (Zhang et al. 2019) are computationally expensive. Furthermore, the large number of covariates in these datasets makes disentangling patientspecific signals from disease and therapy difficult. GeneVector provides a fast and accurate method of cell type classification. We perform cell type classification on a subset of 23,764 cells from the Tumor Immune Cell Atlas (TICA) composed of 181 patients and 18 cancer types (Nieto et al. 2021). The dataset was subset to 2,000 highly variable genes. Cell types were summarized into three main immune cell types: T cells, B/Plasma, and Myeloid cells (**Figure 3A**) from the original annotations (**Figure 3B**). We selected a set of gene markers for each cell type (T cells: CD3D, CD3G, CD3E, TRAC, IL32, CD2; B cells: CD79A, CD79B, MZB1, CD19, BANK1; and Myeloid cells: LYZ, CST3, AIF1, CD68, C1QA, C1QB, C1QC) based on signatures obtained from CellTypist (Conde et al. 2022). For each phenotype, we compute the cosine distance to the normalized expression weighted average of vectors for each gene marker in each cell. The pseudo-probabilities for the three cell types are generated by applying a SoftMax function to the set of cosine distances. The maximum pseudo-probability is used to classify each cell into T cell, B/Plasma, or Myeloid (**Figure 3C**).

**Figure 3:**
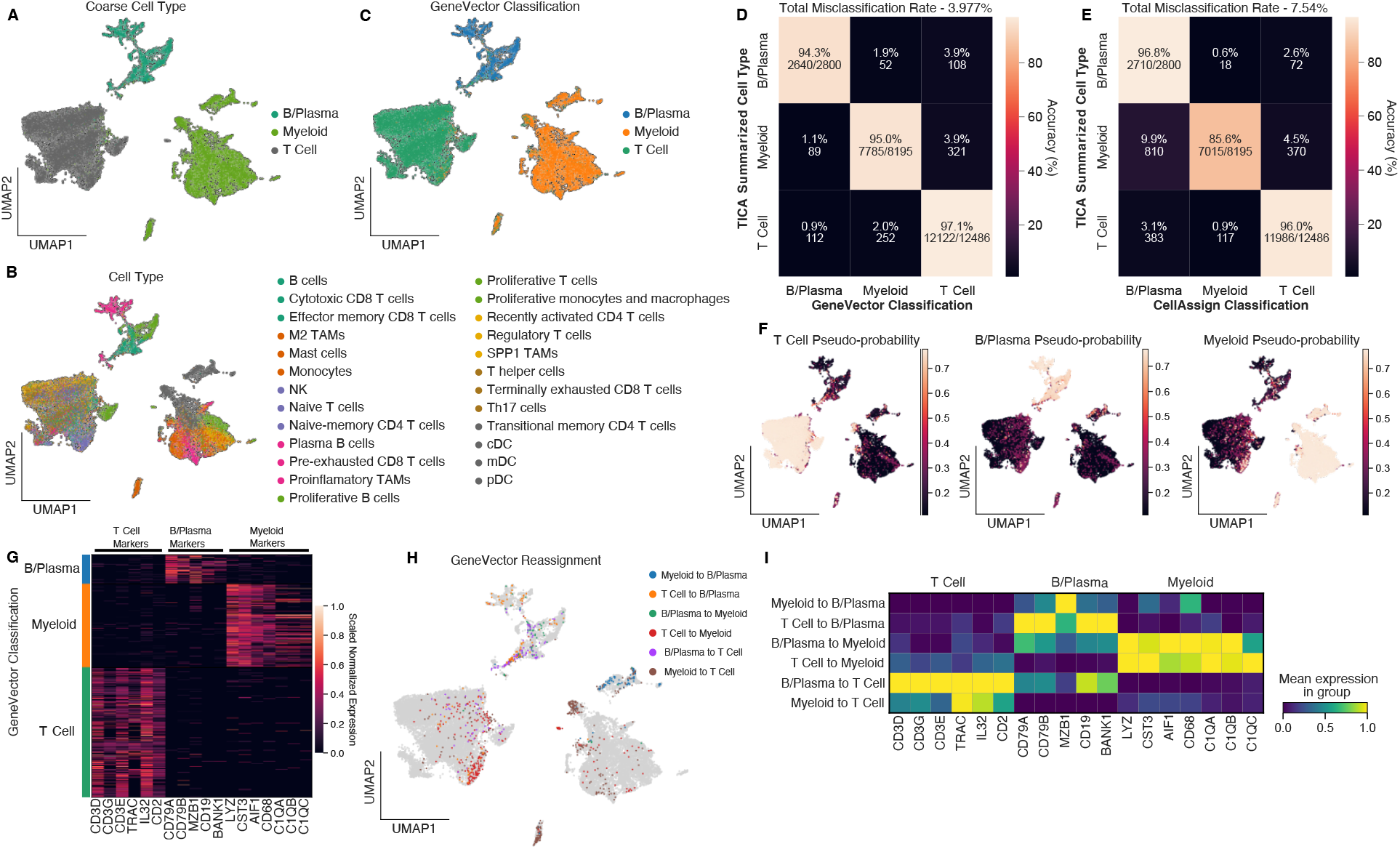
GeneVector Accurately Classifies Cells in TICA Cell Atlas. A) Cells annotated by cell type summarized from the original TICA provided annotations. B) Original annotations provided in TICA. C) GeneVector classification results for each cell type. D) Confusion matrix comparing GeneVector classification with summarized cell types. E) Confusion matrix using the same gene markers with CellAssign shows decreased performance in myeloid cells F) Pseudo-probability values for each summarized cell type. G) Marker gene expression over all cells grouped by GeneVector classification. H) Original annotations reassigned by GeneVector. I) Marker gene mean expression for cell type reassignments highlights misclassifications in B/Plasma cells and doublet transcriptional signatures in T and myeloid cells.

To assess performance, we computed accuracy against the coarse labels in a confusion matrix as the percentage of correctly classified cells over the total number cells for each summarized cell type. We found 97.1% of T cells, 95% of myeloid cells, and 94.3% of B/Plasma cells were correctly classified with respect to the original annotations (**Figure 3D**). Additionally, we classified cells using the same marker genes with CellAssign and found significantly decreased performance in the classification of myeloid cells (85.6%) (**Figure 3E**). Overall, the percentage of cells misclassified using GeneVector (3.977%) showed improvement over CellAssign (7.54%). Using the pseudo-probabilities, GeneVector can highlight cells that share gene signatures including plasmacytoid dendritic cells (pDCs), where cell type definition is difficult (Ziegler-Heitbrock et al. 2023) (**Figure 3F**). For each cell type, we validated that the classified cells are indeed expressing the supplied markers by showing the expression for each marker grouped by classified cell type (**Figure 3G**). For those cells GeneVector reassigned from the original annotations (**Figure 3H**), we examined the mean normalized expression per marker gene and found that many of these cells appear misclassified in original annotations (**Figure 3I**). Cells originally annotated as T cells that were reassigned as B/Plasma by GeneVector show high expression for only B/Plasma markers (CD19, BANK1, CD79A, and CD79B). Additionally, there is evidence that many of these reassigned cells may be doublets. B/Plasma cells reassigned to T or myeloid cells show simultaneous expression of both gene markers. While any computational cell type classification cannot be considered ground truth, cell type assignment with GeneVector is an improvement over CellAssign and demonstrates sensitivity to cells that express overlapping cell type transcriptional signatures.

After learning the gene embedding, GeneVector allows rapid testing of different marker genes and phenotypes in exploratory analysis settings. Increased performance in classification is important given the large variation of markers used to define the same phenotypes across different studies. Cell type prediction can be recomputed interactively within a Jupyter notebook within twenty seconds for even large datasets on most machines (**Supplemental Figure 1**). An additional advantage of having a probability is the ability to map genes from known pathways to a continuous value in each cell. In both phenotype and pathway, demonstration of continuous gradients across cells provides a measure of change and activation that cannot be seen from unsupervised clustering.

### Comparison of Methods in Identification of Interferon Metagenes in 10k Human PBMCs

To identify cell-specific metagenes related to interferon beta stimulation and compare with transcriptional programs identified by PCA and LDVAE loadings, we trained GeneVector using peripheral blood mononuclear cells (PBMCs) scRNA-seq data from 6,855 quality control filtered cells composed of an interferon beta stimulated sample and a control sample (Kang et al. 2018). The original count matrix was subset to 1,000 highly variable genes using the Seurat V3 method (Butler et al. 2018) as implemented in Scanpy (Wolf, Angerer, and Theis 2018). We used previously annotated cell types generated from unsupervised clustering as ground truth labels (T. Stuart et al. 2019). A comparison of the uncorrected UMAP embedding (**Figure 4A**) and subsequent GeneVector-based batch correction (**Figure 4B**) demonstrates correction in the alignment of cell types between the two conditions. However, in contrast to batch correction using Harmony (Korsunsky et al. 2019) (**Figure 4C**), not all variation is lost between the interferon beta stimulated and control cells. Specifically, GeneVector does not align myeloid cell types, suggesting a larger effect of the interferon beta stimulation treatment in these cells.

**Figure 4:**
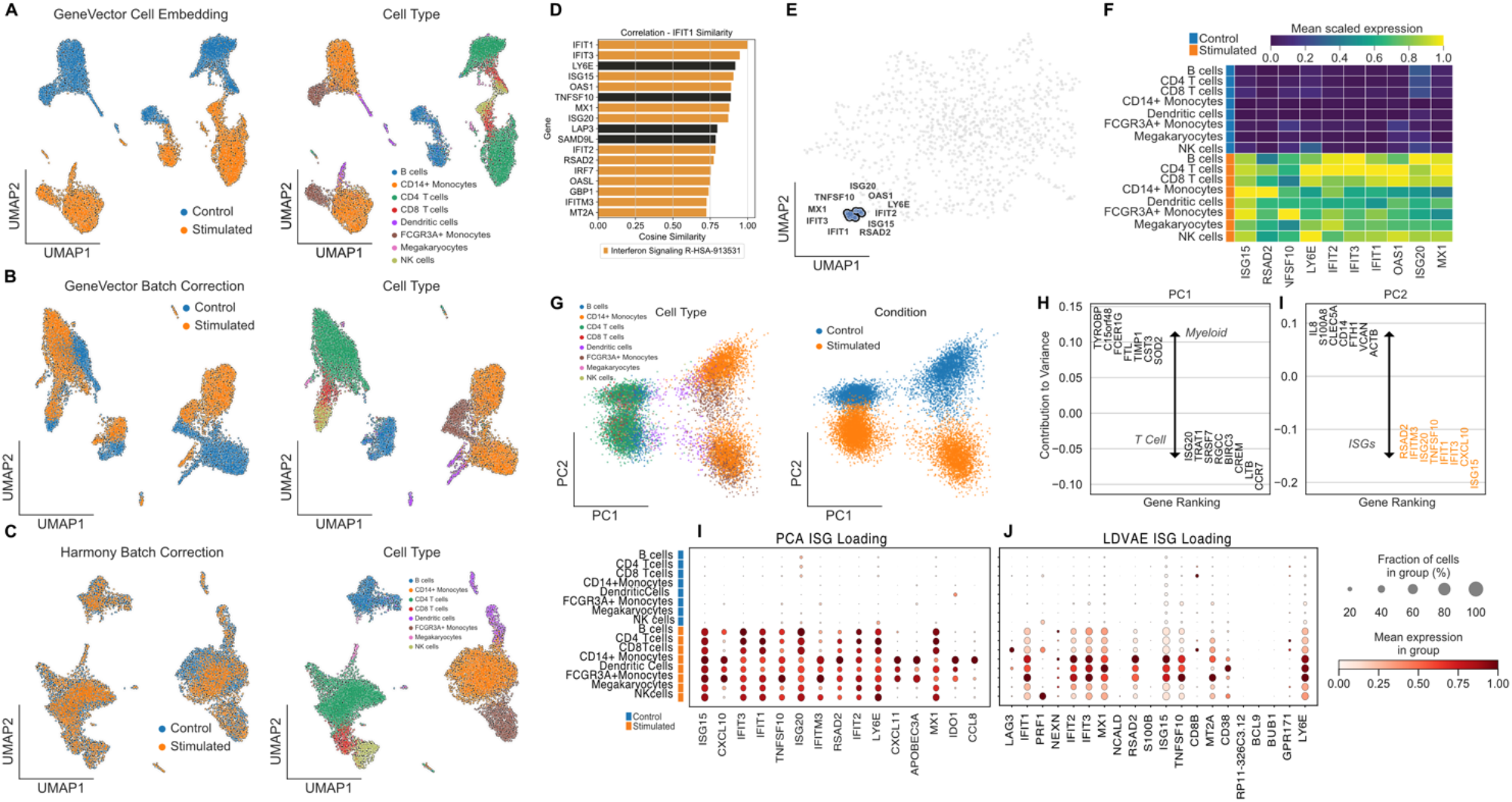
Comparing Methods with Interferon Beta Stimulated 10K PBMCs. A) Uncorrected GeneVector UMAPs showing stimulated condition (left) and cell type annotation (right) on 10k PBMCs with control and interferon beta stimulated cells. B) GeneVector batch corrected UMAPs showing stimulated condition (left) and cell type annotation (right) indicated stronger interferon-beta stimulated response in myeloid cells. C) Harmony batch correction applied to normalized expression eliminates all variation related to interferon beta stimulation. D) Most similar genes by cosine similarity to IFIT1 includes genes found in Reactome interferon related pathways. E) Gene embedding UMAP highlighting an ISG metagene that includes genes most similar to IFIT1. F) Normalized expression of ISG metagene shows increased expression across stimulated cells without cell type specific effects. G) PCA embedding colored by cell type (left) and stimulation (right). H) Top genes by contribution to variance indicates PC1 defines cell type. I) Top genes by contribution to PC2 defined by ISG stimulation. J) Normalized expression of ISG related loadings from PCA (left) and LDVAE (right) includes cell type specific markers intermixed with ISGs.

As a method of both validation and exploration, GeneVector provides the ability to query similarity in genes. For a given target gene, a list of the closest genes sorted by cosine similarity can be generated. This is useful in both validating known markers and identifying the function of unfamiliar genes by context. As previously described, the genes most similar to IFIT1 (**Figure 4D**) include a large proportion of genes found in the Reactome pathway Interferon Signaling (R-HSA-913531) (Gillespie et al. 2022). After clustering gene vectors, we identify a single metagene that includes these genes (IFIT1, IFIT2, IFIT3, ISG15, ISG20, TFGS10, RSAD2, LYSE, OAS1, and MX1). The ISG metagene can be visualized on a UMAP generated from the gene embedding, like the familiar cell-based visualizations common in scRNA-seq studies (**Figure 4E**). The mean expression of each gene identified in the ISG metagene is significantly higher in interferon beta stimulated cells over control cells (**Figure 4F**). Importantly, the increased expression is found in each cell type, indicating a global relation to treatment.

To compare the ISG metagene with results generated from PCA loadings, we performed PCA on the normalized gene-by-cell expression matrix (**Figure 4G**) and colored the embedding by cell type and treatment. After computing the PCA loadings using Scanpy (Wolf et al. 2018), we identified the top genes by contribution score to variation in the first and second principal components. The first principal component (PC1) explains variation related to cell type and the differences between myeloid (TROBP, FCER1G, FTL, CST3) and T cells (LTB, CCR7) (**Figure 4H**). The second principal highlights the variation related to interferon beta stimulation and includes the genes found in the ISG metagene generated by GeneVector (**Figure 4I**). However, the increased effect of interferon stimulation in myeloid cells, conflates myeloid specific ISGs with the interferon signature. One such gene is CXCL10, which shows cell type specificity to myeloid cells (**Figure 4J**) and is not found in the interferon signaling Reactome pathways. Additionally, IFITM3 shows increased expression only in myeloid cells within these PBMCs. In contrast, GeneVector produces a metagene that groups myeloid specific genes into a single metagene including CXCL10 and IFITM3. A full list of metagenes produced by GeneVector is presented in **Supplemental Figure 2**. Among these metagenes, we identify transcriptional programs specific to each cell type and treatment condition, including those found in the least represented cell type Megakaryocytes (132 of 14,038 cells).

To compare the GeneVector ISG metagene with LDVAE, we trained an LDVAE model using 10 latent dimensions for 250 epochs with control and stimulated batch labels in the SCVI framework. In contrast to the specificity of the GeneVector ISG metagene that includes only interferon stimulated genes, the nearest LDVAE loading mixes interferon-related genes with markers of T cell activation (PRF1) and T cell dysfunction (LAG3) (**Figure 4K**). With respect to PCA and LDVAE loadings, GeneVector identified an ISG metagene that is not confounded by cell type and includes only interferon pathway related genes.

### Metagenes Changes Between Primary and Metastatic Site in HGSOC

Studies with scRNA-seq data sampled from multiple tumor sites in the same patient provide a wide picture of cancer progression and spread. As these datasets grow larger and more complex, understanding the transcriptional changes that occur from primary to metastatic sites can help identify mechanisms that aid in the process of the invasion-metastasis cascade. GeneVector provides a framework for asking such questions in the form of latent space arithmetic. By defining the difference between two sites as a vector, where the direction defines transcriptional change, we identify metagenes associated with expression loss and gain between primary and metastasis sites from six patients in the Memorial Sloan Kettering Cancer Center SPECTRUM cohort of patients with high-grade serous ovarian cancer (HGSOC) (Vázquez-García et al. 2022).

A set of 270,833 cells quality control filtered cells from adnexa (primary) and bowel (metastasis) samples were processed with GeneVector (**Figure 5A**). The dataset was subset to 2,000 highly variable genes and cells were classified to one of six cell types using gene markers curated for HGSOC and two markers for cancer cells (EPCAM and CD24) (Vázquez-García et al. 2022). We compared the classification accuracy to the original annotations generated from CellAssign for each cell type in a confusion matrix (**Figure 5B**). Accuracy reached 99.7% in three cell types with a minimum classification rate of 94.1%. In cells where GeneVector annotated differently than previous annotations, there is evidence from the differentially expressed genes that these cells may have been initially mislabeled. GeneVector reassigned a subset of cancer cells to fibroblast and the differentially expressed genes between these cells and the cells annotated as cancer by GeneVector highlighted fibroblast cell type markers including COL1A1 (**Supplemental Figure 3A**). In B/Plasma cells reassigned as T cells, the differentially expressed genes highlight B cell receptor genes IGK and IGLC2 (**Supplemental Figure 3B**). Finally, in B/Plasma cells classified as myeloid, the top differentially expressed genes include known canonical markers for macrophage/monocyte cells (TYROBP, LYZ, CD4, and AIF1) (Conde et al. 2022) (**Supplemental Figure 3C**).

**Figure 5:**
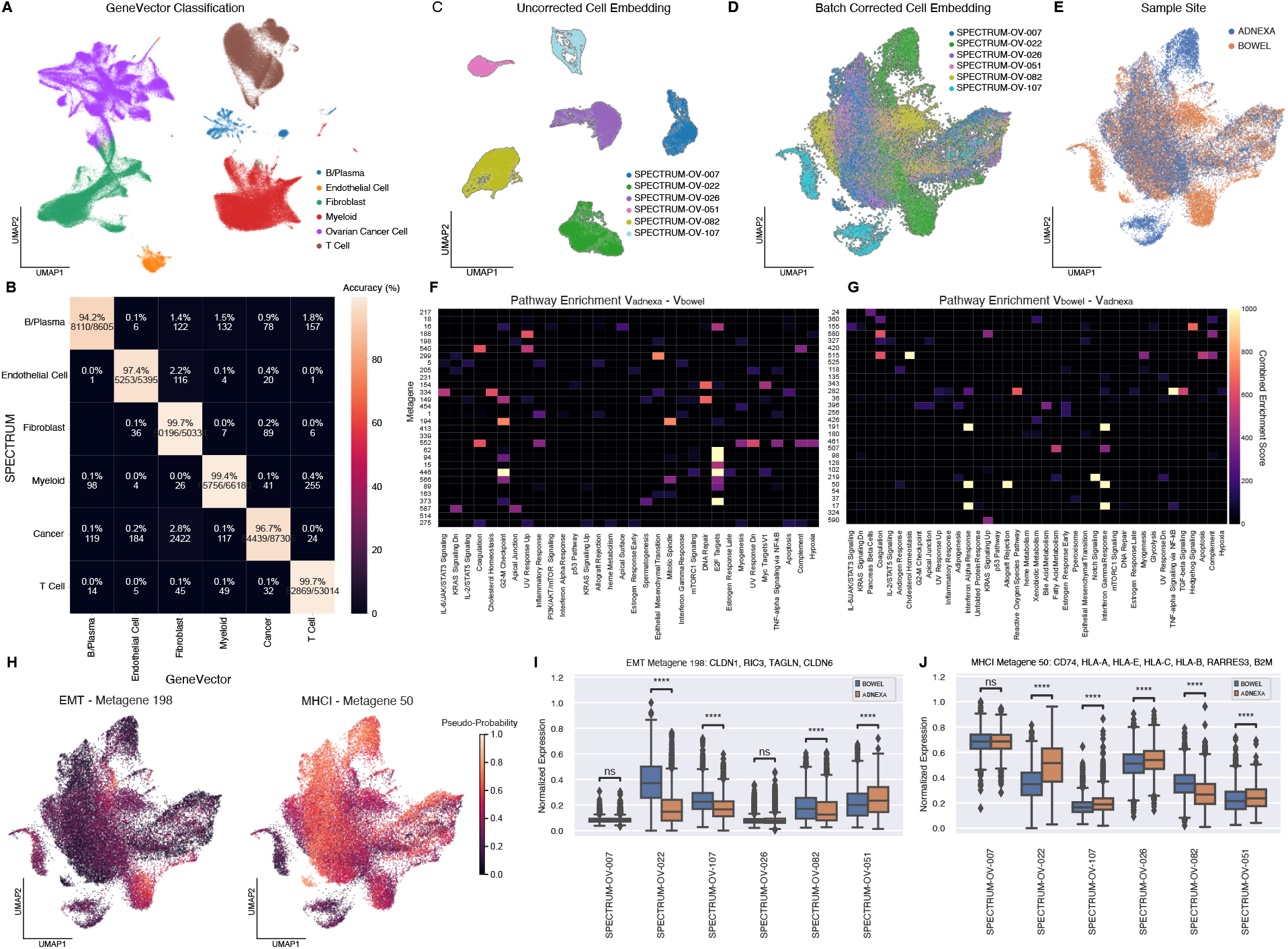
Metagenes associated with directional difference in HGSOC cancer cells from adnexa to bowel. A) UMAP of HGSOC cells with classified by GeneVector. B) Confusion matrix of accuracy comparing SPECTRUM annotated cell types with GeneVector classification. C) Uncorrected UMAP of cancer cells from patients with adnexa and bowel samples. D) GeneVector batch corrected UMAP with patient labels. E) Site labels on batch corrected UMAP. F) Pathway enrichment for top 30 metagenes by cosine similarity to V_adnexa_ – V_bowel_. G) Hallmark pathway enrichment for top 30 metagenes by cosine similarity to V_bowel_ – V_adnexa_ H) Pseudo-probabilities for metagenes associated with up-regulation in bowel to adnexa (Epithelial-to-Mesenchymal Transition) and down-regulation (MHC Class I). I) EMT metagene significantly up regulated in four of six patients by gene module score. J) MHCI metagene significantly downregulated in the metastatic site (bowel) in three of six patients by gene module score. (**** p < 0.0001, ns=not significant)

We recomputed the gene embedding and metagenes on only those cells classified as cancer by GeneVector with both adnexa and bowel samples. We found these cells exhibited large patient specific batch effects (**Figure 5C**) and applied GeneVector batch correction (**Figure 5D-E**). To understand the changes between primary and metastatic sites, we computed an average vector for all cells from the adnexa (v_adnexa_) as the primary site and bowel (v_bowel_) as the site of metastasis. We mapped the top 30 most similar metagenes to vectors representing expression gain in metastasis (v_adnexa_ - v_bowel_) (**Figure 5F**) and expression loss (v_bowel_ - v_adnexa_) (**Figure 5G**). Gene Set Enrichment Analysis (GSEA) using GSEAPY with Hallmark gene set annotations from Enrichr (Kuleshov et al. 2016) was performed to assess whether metagenes were enriched for genes from known pathways. Metagenes enriched for E2F targets and Epithelial-to-Mesenchymal Transition (EMT) pathway genes were found gained in metastasis. Conversely, the set of metagenes representative of loss from adnexa to bowel included MHC Class I (HLA-A, HLA-B, HLA-C, HLA-E, and HLA-F) and the transcriptional regulator B2M, suggesting a means of immune escape via loss of MHC Class I expression and higher immune pressure in metastatic sites may increase the potential fitness benefit of MHC Class I loss. For both the EMT and MHCI metagenes, pseudo-probabilities computed using GeneVector highlight pathway activity in either site in the UMAP embedding (**Figure 5H)**. Using gene module scoring (Satija et al. 2015), we examined the change between sites in each patient for the EMT and MHCI metagenes. We found that the MHCI metagene is significantly downregulated in metastatic sites in four of six patients (**Figure 5I**). Conversely, the EMT metagene was significantly up regulated in metastatic sites for three of six patients (**Figure 5J**). The ability to phrase questions about transcriptional change as vector arithmetic provides a powerful platform for more complex queries than can be performed with differential expression analysis alone.

### Metagenes Associated with Cisplatin Treatment Resistance

Understanding the transcriptional processes that generate resistance to chemotherapies has immense clinical value. However, the transcriptional organization of resistance is complex with many parallel mechanisms contributing to cancer cell survival (Shen et al. 2012). We analyzed longitudinal single cell RNA-seq collected from a triple negative breast cancer patient-derived xenograft model (SA609 PDX) along a treated and untreated time series (Salehi et al. 2021). Using a total of 19,799 cancer cells with treatment and timepoint labels (**Figure 6A**), we generated metagenes to identify programs potentially related to cisplatin resistance. For each metagene, we computed gene module scores (Satija et al. 2015) over the four timepoint (X1, X2, X3, and X4) within the treated and untreated cells. Using these scores, we calculated linear regression coefficients over the four time points and selected candidate chemotherapy resistant and untreated metagenes (β_*treated*_ and β_*untreated*_) over a coefficient threshold (β_*treated*_ >0.1 and adj-p < 0.001) with an untreated coefficient less than the treated coefficient (β_*treated*_ > β_*untreated*_) **(Figure 6B**). Mean expression was computed for each timepoint in five metagenes that were identified as treatment specific (**Figure 6C**).

**Figure 6:**
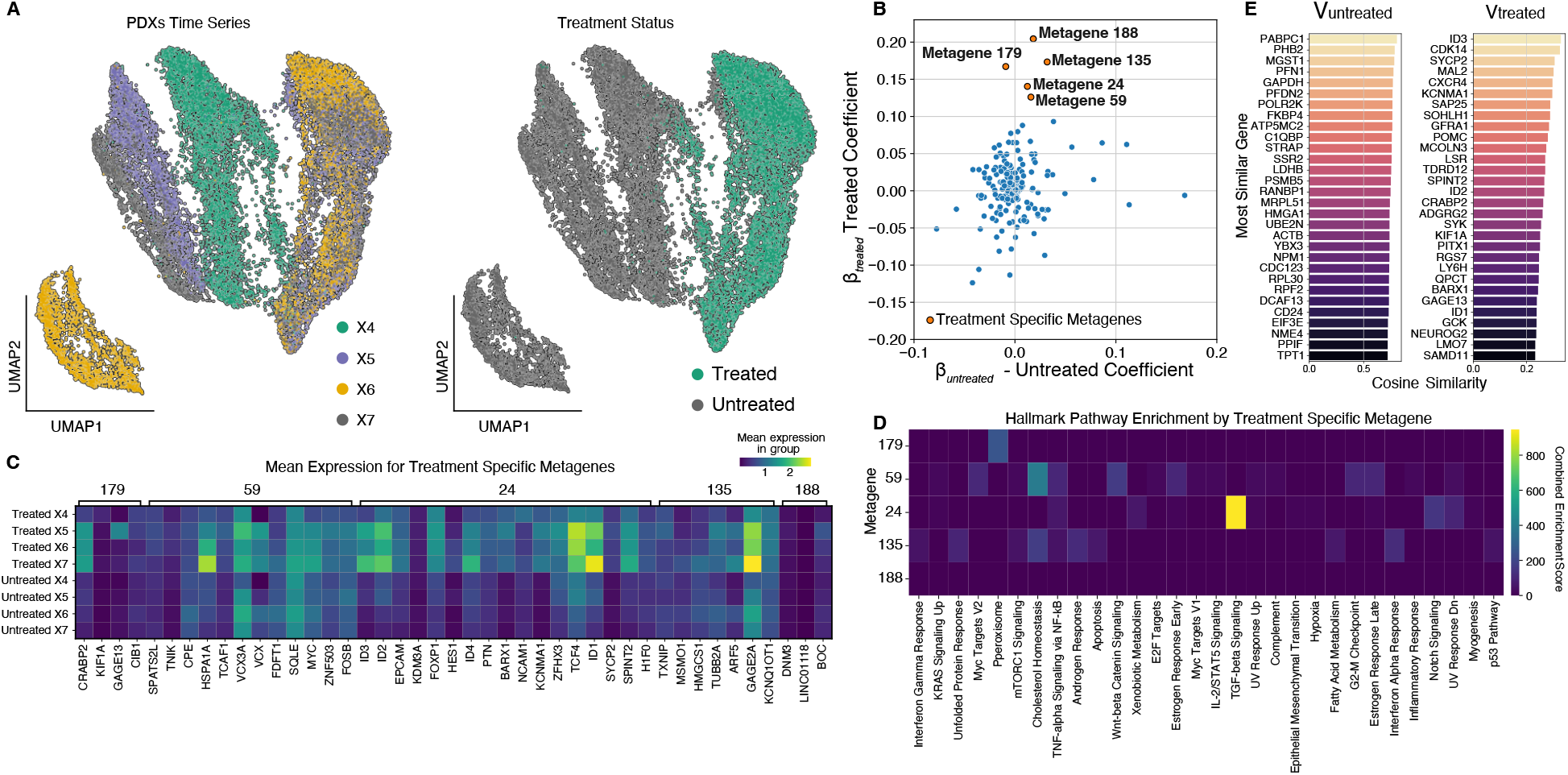
Analysis of Metagenes in Cisplatin-treated PDX Time Series. A) UMAP of cisplatin treated PDX cells annotated by time point (left) and treatment status (right). B) Metagenes plotted with respect to regression coefficients (β_treated_ and β_untreated_) over four timepoints in either treated or untreated cells with Bonferroni adjusted p-values < 0.001. C) Hallmark combined enrichment scores for candidate chemo resistant metagenes. D) Gene expression profiles for each timepoint for the five metagenes associated with increase in treatment. Metagene 24 is associated with TGF-beta signaling includes EPCAM, FOXP1, and ID family genes with known cisplatin resistance function. E) Most similar genes for treated and untreated cells computed from cosine similarity to global vectors.

Pathway enrichment on each of the five metagenes using GSEAPY with Hallmark gene set annotations from Enrichr (Kuleshov et al. 2016) showed that metagene 24 was enriched for TGF-beta signaling (**Figure 6D**), a pathway frequently up-regulated during chemo-resistance (Bhola et al. 2013). This metagene includes genes ID1, ID2, ID3, ID4, EPCAM, and FOXP1, all of which have been found to be overexpressed in chemotherapy-resistant samples in several cancer types (Zhang et al. 2006; Yamano et al. 2010; Roberts et al. 2005). GeneVector also identified these genes in global expression differences between treated and untreated cells from the set of most similar genes to vectors V_treated_ and V_untreated_ (**Figure 6E**). Several studies have implicated multiple resistance mechanisms involving FOXP1 including transcriptional regulation, immune response, and MAPK signaling (Zhu et al. 2015; Choi et al. 2016; Hu et al. 2015). Additionally, GeneVector identifies EPCAM, whose high expression has been associated with increased viability of cancer cells in diverse cancer types (Sun et al. 2018; Imrich, Hachmeister, and Gires 2012). EPCAM has been shown to have a role in resistance to chemotherapy in both breast and ovarian cancers through WNT signaling and Epithelial-Mesenchymal Transition (EMT) (Tayama et al. 2017; Latifi et al. 2011).

## Discussion

In this paper we propose GeneVector, a method for building a latent vector representation of scRNA expression capturing the relevant statistical dependencies between genes. By borrowing expression signal across genes, GeneVector overcomes sparsity and produces an information dense representation of each gene. The resulting vectors can be used to generate a gene co-expression graph, and can be clustered to predict transcriptional programs, or metagenes, in an unsupervised fashion. Metagenes can be related to a cell embedding to identify transcriptional changes related to conditional labels or time points. We show that gene vectors can be used to annotate cells with a pseudo-probability, and that these labels are accurate with respect to previously defined cell types.

We show that a single GeneVector embedding can be used for many important downstream analyses. In interferon-stimulated PBMCs, we identify a cell type independent ISG metagene that summarizes interferon-stimulation across cell types and is not conflated with cell type signature. We demonstrate accurate cell type assignment across 18 different cancers in 181 patients described in the Tumor Immune Cell Atlas (TICA). In high grade serous ovarian cancer, we identify metagenes that describe transcriptional changes from primary to metastatic sites. Our results implicate the loss of MHC class I gene expression as a potential immune escape mechanism in ovarian cancer metastasis. In cisplatin treated TNBC PDXs, GeneVector uncovers transcriptional signatures active in drug resistance, most notably metagenes enriched in TGF-beta signaling. This signaling pathway is a cornerstone in cancer progression since it promotes EMT transition and invasion in advanced cancers; it is the target of various therapies, but success has been mixed (M. Zhang et al. 2021) making it even more important to employ tools that identify the multitude of players contributing to therapy response.

Identifying correlations across scRNA data is a fundamental analysis task, necessary for identifying cells with similar phenotype or activity, or genes with similar pathways or functional relationships. As has been shown by us and in previous work (Margolin et al. 2006; Chan, Stumpf, and Babtie 2017; Xu et al. 2022), the sparsity and non-linearity of scRNA data impact the performance of both standard measures of correlations between variables and global analysis of assumed linear correlations using PCA. While some methods tailor complex custom probabilistic models to the specific properties of scRNA data, GeneVector instead builds upon MI, a simple yet powerful tool for calculating the amount of information shared between two variables. We show that MI and the vector space trained from the MI matrix both capture relevant gene pair relationships including between TF activators and repressors and their targets. Nevertheless, GeneVector is unable to discern repression from activation, as it builds MI that is agnostic to the direction of the statistical dependency. Pearson correlation, theoretically sensitive to the direction of a dependency, also performs poorly. Due to the high-level of sparsity, absence of expression is not a significant event and repressed expression could just as easily be explained by under-sampling an expressed gene. We suggest that identifying negative regulation and mutually exclusive expression is one of the more difficult problems in scRNA-seq analysis.

As shown with correlation, the objective function employed in training GeneVector has a significant effect on the resulting gene embedding. Mutual information calculated empirically from the histogram of binned expression counts for gene pairs is limited by the available number of cells, fidelity of the counts, and discretization strategy. By modeling the underlying distribution for each gene more accurately, the joint probability distribution between genes can more accurately reflect expression-based relationships and improve model results. Additionally, while only a one-time cost, the MI calculation is computationally expensive. By improving the calculation of mutual information, others have achieved improved performance in related tasks including the identification of GRNs (Lachmann et al. 2016).

The high dimensionality of scRNA and the vast complexity of biological systems to which it is applied necessitate analytical tools that facilitate intuitive and efficient data exploration and produce easily interpretable results. GeneVector performs upfront computation of a meaningful low dimensional representation, transforming sparse and correlated expression measurements into a concise vector space summarizing the underlying structure in the data. The resulting vector space is amenable to intuitive vector arithmetic operations that can be composed into higher level analyses including cell type classification, treatment related gene signature discovery, and identification of functionally related genes. Importantly, the vector arithmetic operations and higher-level analyses can be performed interactively, allowing for faster iteration in developing cell type and context specific gene signatures or testing hypotheses related to experimental covariates. GeneVector is implemented as a python package available on GitHub (https://github.com/nceglia/genevector) and installable via PIP.

## Online Methods

### Gene Expression Mutual Information

In NLP applications, vector space models are trained by defining an association between words that appear in the same context. In single cell RNA sequencing data, we can redefine this textual context as co-expression within a given cell and mutual information across cells. The simplest metric to define association is the overall number of co-expression events between genes. However, the expression profiles over cells may differ due to both technical and biological factors. To summarize the variability in this relationship, we generate a joint probability distribution on the co-occurrence of read counts. The ranges of each bin are defined separately for each gene based on a user defined number of quantiles. By defining the bin ranges separately, the lowest counts in one gene can be compared directly to the lowest counts in another gene without need for further normalization. Using the joint probability distribution, we compute the mutual information between genes defined in **Equation 1**. The mutual information value is subsequently used as the target in training the model, allowing us to highlight the relationship between genes independent of normalization methods as a simple, single-valued quantity.

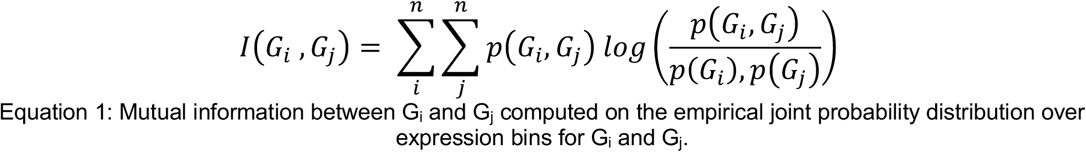

### Model Training

A neural network is constructed from a single hidden layer corresponding to the size of the latent space vectors. A set of independently updated weights connects the one-hot encoded input and output layers that are defined from each pair of genes. These weights, *w_1_* and *w_2_*, are matrices with dimensions equal to *N* expressed genes by *I* hidden units. Initial values for *w_1_* and *w_2_* are generated uniformly on the interval −1 to 1. The objective function, as a minimization of least squares, is defined in **Equation 2**. The final latent space, defined as the gene embedding, is computed as the vector average of weights *w_1_* and *w_2_.* For NLP applications, this is a preferred approach over selecting either weight matrix (Pennington, Socher, and Manning 2014). Each co-expressed gene pair is used as a single training example. The maximum number of examples for a full training epoch is given by the total number of co-expressed gene pairs. Weights are batch updated with adaptive gradient descent (ADADELTA) (Duchi et al. 2011). Training is halted at either a maximum number of epochs, or when the change in loss falls below a specified threshold. The model is implemented in PyTorch.

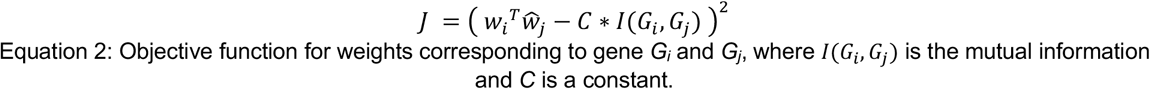

### Gene Vectors

The cosine similarity function, defined in **Equation 3**, is used to measure similarity between vectors, defined as 1 - cosine distance or dot product. Values closer to 1 indicate strong association within the dataset. A gene vector is defined as the learned weights in the gene embedding for a particular gene. Vectors describing groups of cells are generated by computing a weighted average vector, described in **Equation 4**, from a set of individual gene vectors.

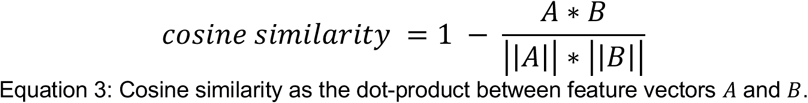

To assign a vector to each cell, the average vector is computed across the gene embedding weighted by counts observed in each cell. The matrix of all cell vectors in the dataset is defined as the cell embedding. This cell embedding can be used in place of PCA or embeddings obtained with variational auto-encoders.

Each cell vector maintains a linear relationship with the gene embedding. The computation of vectors describing groups of cells is described in Equation 4.

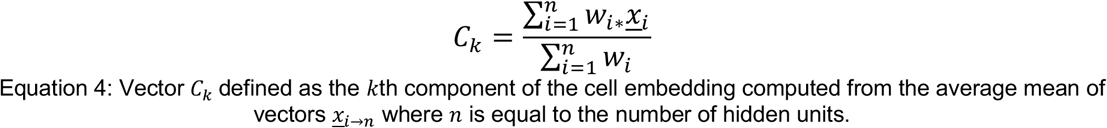

### Co-expression Similarity Graph

A co-expression similarity graph is constructed from cosine similarity between each pair of genes using Scanpy *neighborhood* function with a default value for *k* of 15. A node in the graph represents a single gene and edges are weighted by cosine similarity. To generate metagenes, we apply Leiden clustering to the neighborhood graph with a user defined resolution argument. We have found that metagene clustering is robust to a range of resolution values ranging from 15 to 1000 when assessing the identification of an MHCI-like metagene in a single patient in the SPECTRUM cohort (**Supplemental Figure 4**).

### Cell Type Assignment

A set of cell types with user defined marker genes are used to perform a pseudo-probabilistic phenotype assignment in a single cell. A representative vector for the cell type is computed from the user defined marker genes weighted by the normalized expression of the cell to be classified. The cosine distances of each cell type vector with cell vector are passed through a SoftMax function, given in **Equation 5**, to provide a pseudo-probability distribution for each phenotype. The argument maximum of this distribution is used to classify the most likely cell type for a given cell. This procedure is repeated for every cell in the dataset.

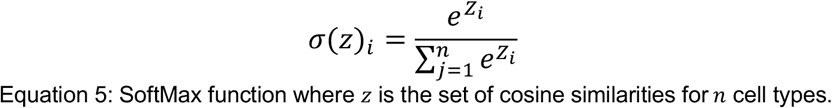

### Generation of Predictive Genes

The method of phenotype assignment can be reversed to produce a set of genes that are most similar, or predictive, to any group of cells. The cell vectors belonging to this group can be averaged to generate a group vector representing a label in the dataset. The transcriptional signature specifically associated with this group of cells can be computed using vector arithmetic V_group_ – V_dataset_, where V_dataset_ is the average vector over all cells. Gene vectors are sorted by cosine similarity to the group vector to produce a ranked list of candidate genes that are most like the set of cells in the group.

### Batch Correction

Batch correction is applied to cells over a given set of batch labels with the goal of correcting the difference between these batches and a user-specified reference batch. With the set of cells labeled for a batch, we can compute an average batch vector. A correction vector can then be computed for each batch using vector arithmetic to describe the direction and magnitude of the change from the batch to the reference. The correction vector is then added to each cell in the batch. After computing the cell embedding, the following procedure is applied to each batch label:

1. Compute the average vector *V_batch_* for a set of cells.
2. Compute the reference vector *V_reference_* for cells in the reference batch.
3. Compute a correction vector *V_correction_* = *V_reference_* – *V_batch_.*
4. For each cell in the batch, subtract *V_correction_* from each cell vector.
5. Repeat for all *n* batches.

### Time-series Analysis of Metagenes

Linear regression is applied to gene module scores (Satija et al. 2015) over a series of time points for each metagene resulting in a p-value and slope coefficient from each series. Metagenes with a Bonferroni adjusted p-value < 0.001 and a treated coefficient greater than the untreated coefficient, with a minimum threshold of 0.1 were selected as treatment specific gene programs.

## DATA AVAILIBILITY

*The authors declare that the data needed to reproduce analysis in this study are available to download from Google drive using example Jupyter notebooks provided in the GitHub repository (https://github.com/nceglia/genevector/example).*

## CODE AVAILIBILITY

*The authors declare that the code needed to reproduce analysis and findings of this study are available in the GitHub repository (https://github.com/nceglia/genevector) and the GeneVector python package can be installed directly from using the Python installer package (PIP).*

## AUTHOR CONTRIBUTIONS

*N.C. developed method, implemented software, performed analysis, authored manuscript. Z. S. developed method, guided analysis framework, and authored manuscript. S.S.F. developed method, performed data analysis, and authored manuscript. F.U. performed analysis. V.B. performed data analysis and developed visualizations. N.R. authored manuscript and provided analysis guidance. B.B. provided analysis guidance, biological interpretations, and manuscript editing. A.C. provided analysis guidance, biological interpretations, and manuscript editing. S.S. provided biological interpretations and manuscript editing. F.K. provided biological interpretations and manuscript editing. S.A. provided biological interpretations. B.G. supervised method development. S.P.S. supervised method development. A. M. developed method, authored manuscript, and supervised method development.*

## Supplemental Figures

**Supplemental Figure 1:**
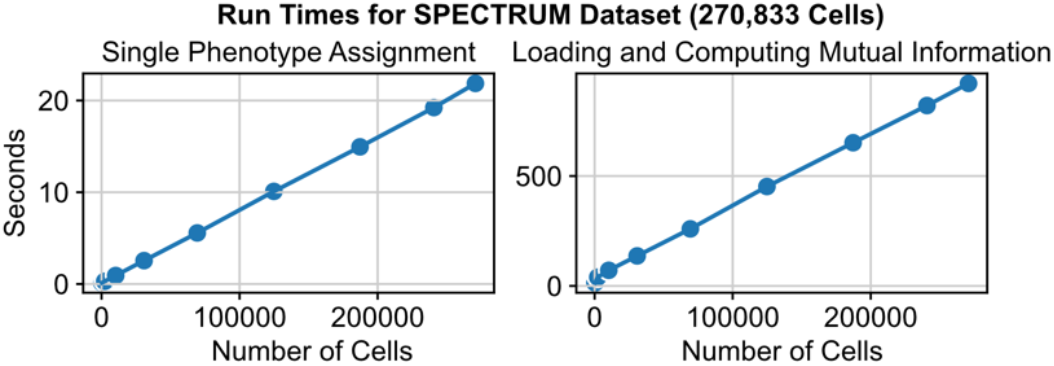
Run times for increasing number of cells for assigning a phenotype to each cell using five gene markers and the time needed to compute the mutual information for 1,000 genes. Both operations scale linearly with input size. Each time was computed on a subsample of the SPECTRUM dataset.

**Supplemental Figure 2:**
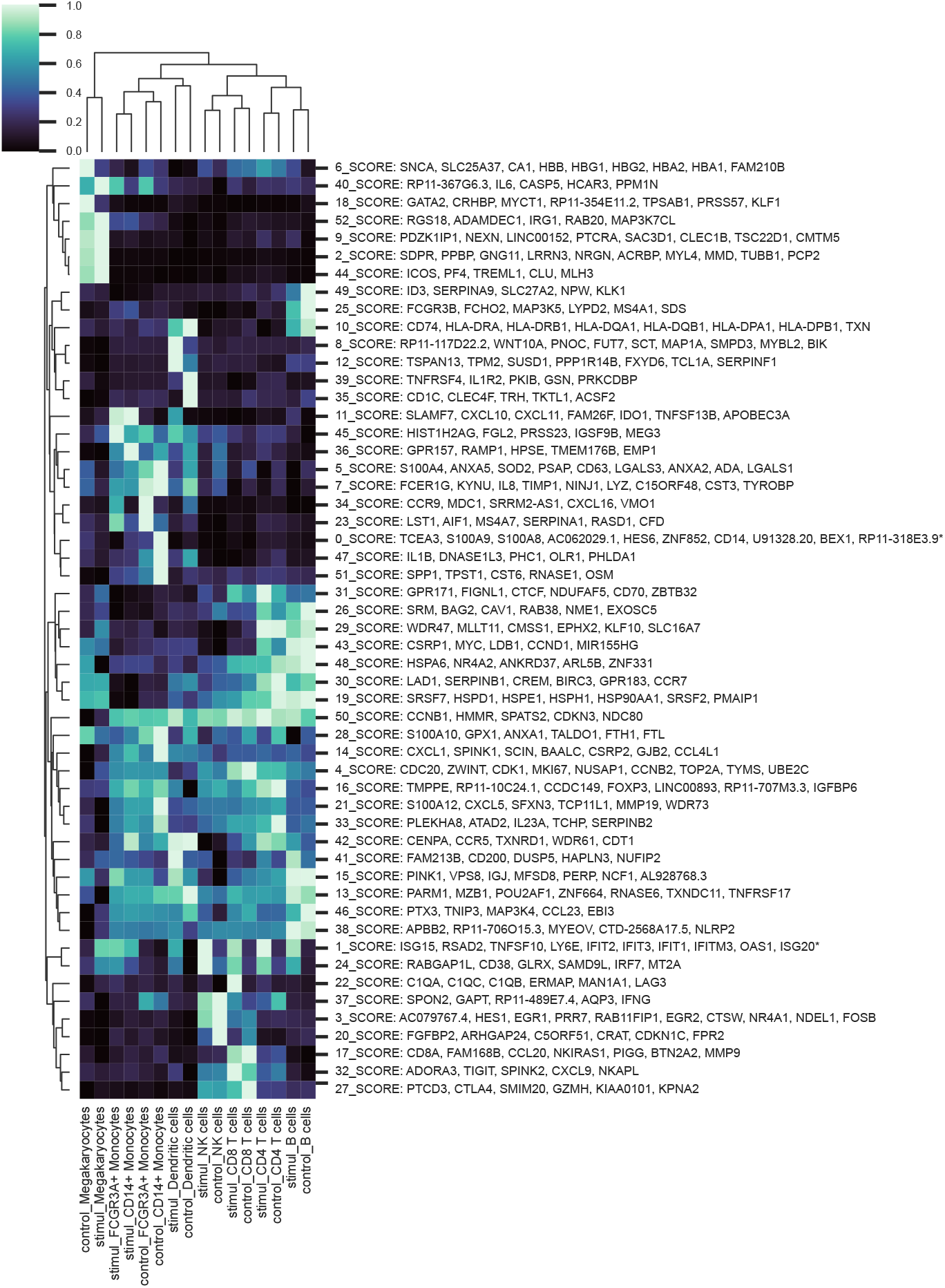
Gene module scores for all metagenes identified in inteferon stimulated PBMCs with greater than three genes. * Indicates additional gene membership. Metagenes programs can be found for each individual cell type and treatment condition including very small cell populations like Megakaryocytes (132 of 14,038 cells).

**Supplemental Figure 3:**
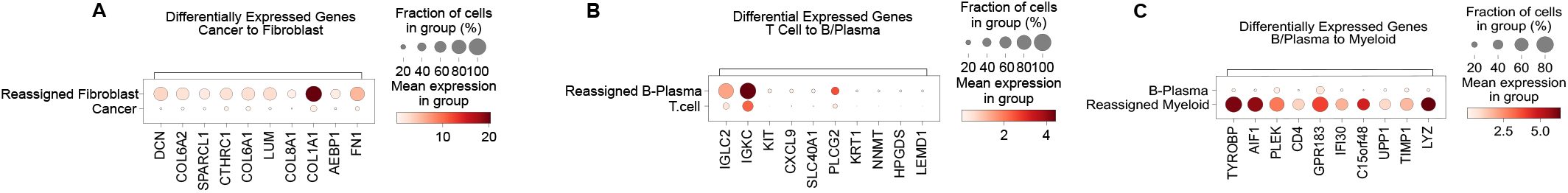
A) Differentially expressed genes in cells classified as fibroblast cells that were classified as cancer using CellAssign highlights the COL1A1. B) Differentially expressed genes in cells classified as B/Plasma cells that were classified as T cells using CellAssign highlights the genes IGKC and IGLC2. C) Differentially expressed genes in cells classified as Myeloid cells that were classified as B/Plasma using CellAssign highlights the genes TYROBP, AIF1, CD4, and LYZ.

**Supplemental Figure 4:**
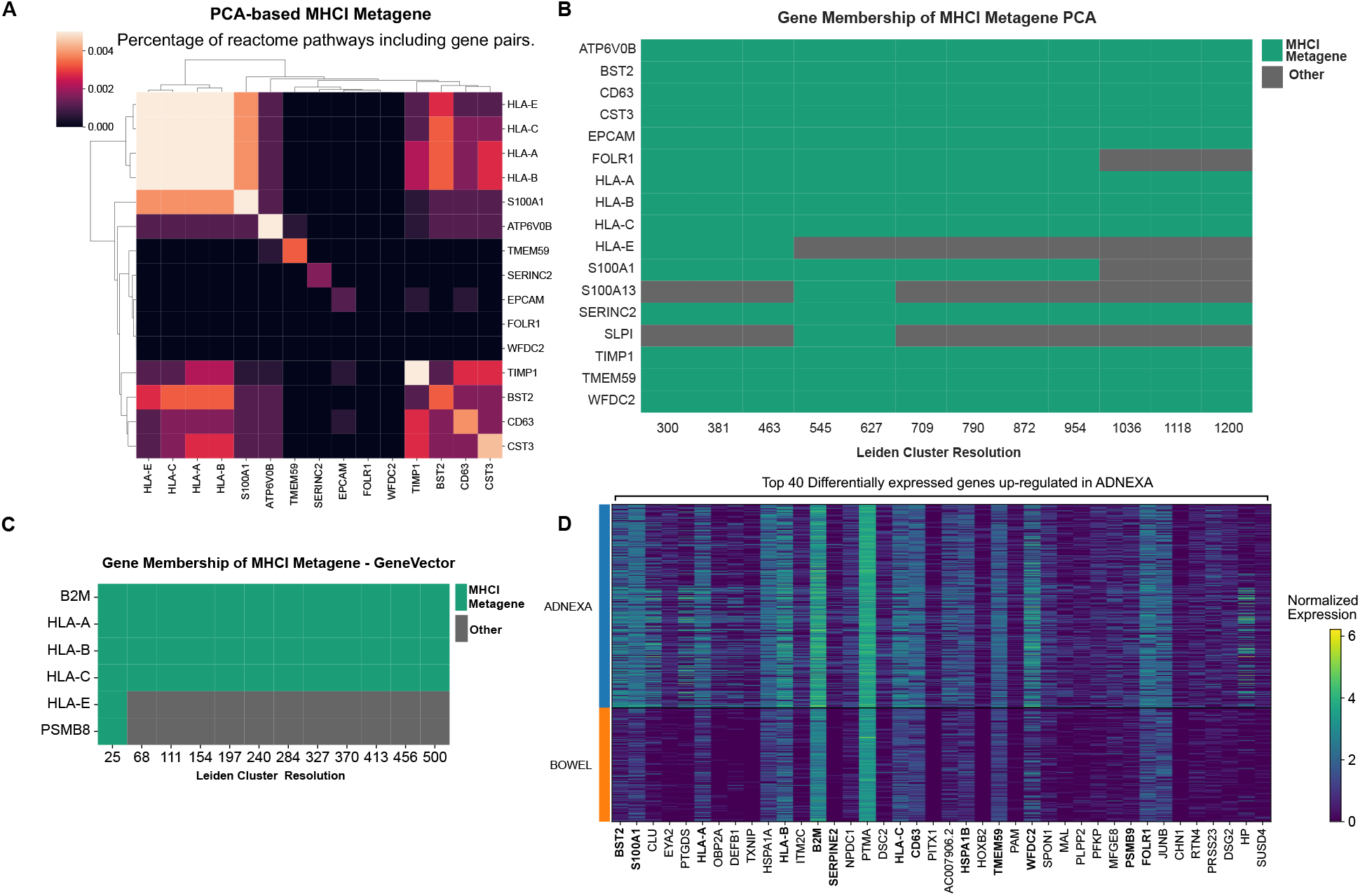
A) Percentage of Reactome pathways where gene pairs from the PCA-based MHCI metagene were found together. B) Gene membership in MHCI metagene over multiple Leiden resolution values highlights robust clustering of HLA genes and non-pathway specific genes. C) Gene membership of MHCI metagene generated from GeneVector robustly selects HLA specific genes. D) Differentially expressed genes up regulated in adnexa includes non-specific pathway genes alongside MHCI genes.

